# Habitat fragmentation compromises the population dynamic of the globally near-threatened Straight-billed Reedhaunter (*Limnoctites rectirostris*)

**DOI:** 10.1101/468488

**Authors:** Maycon S. S. Gonçalves, Priscila S. Pons, Felipe C. Bonow, Vinicius A. G. Bastazini, José A. Gil-Delgado, Germán M. López-Iborra

## Abstract

Understanding the consequences of habitat fragmentation to biological populations is crucial to develop sound conservation polices. The Straight-billed Reedhaunter (*Limnoctites rectirostris*) is a little known and threatened Passeriform that is highly dependent Erygo wetlands patches. Here, we evaluated the effects of habitat fragmentation on populations of the Straight-billed Reedhaunter, during the construction of a water reservoir in southern Brazil. During eight months, we monitored five Eryngo wetlands patches occupied (n=3) and no occupied (n=2) by Straight-billed Reedhaunter individuals, collecting data on their temporal occupancy patterns and registering new fragmentation events in formally continuous habitat patches. We evaluated the consequences of habitat fragmentation on the probabilities of patch occupancy, colonization and extinction of populations of the Straight-billed Reedhaunter using an information-theoretic approach. Out of the three patches occupied by Straight-billed Reedhaunter, two were not altered by construction activities and their populations were present during the entire study period. After fragmentation events, local extinction in one of the wetland patches was observed, and individuals were sporadically observed in two other initially unoccupied sites. The model in which fragmentation affected only the extinction probability was the most plausible among the set of candidate models. Fragmentation greatly increased the chance of local population extinction within patches. Our results indicate that the conservation of populations of the Straight-billed Reedhaunter is highly dependent on continuous and unaltered wetland patches.

## Introduction

Continental wetlands are among the most threatened ecological systems in the World (Davidson 2014). In South America, habitat fragmentation of some regions has reduced the surface of this ecosystem to less than 10% of its original area (Maltchik et al. 2003; Guadagnin et al. 2005). One of the most interesting and little-known continental aquatic systems in South America refers to those dominated by Eryngo *Eryngium pandanifolium* Cham. & Schltdl. (Apiaceae), locally known as *gravatazais*. These wetlands are distributed in the form of patches in the grassland landscape and are periodically altered or destructed by many anthropic disturbances, such as agriculture, intensive livestock, intentional fires and drainage for dam construction (Irgang 1999; López-Lanús et al. 1999; Bencke et al. 2003; Volcan et al. 2014).

Collected for the first time in June 1833 by Charles Darwin (Steinheimer 2004), the Straight-billed Reedhaunter (*Limnoctites rectirostris*) is a South American aquatic passerine that strongly depends on the wetlands dominated by Eryngo (BirdLife International 2016). The Straight-billed Reedhaunter has no known migratory movements and its geographical distribution is limited to southern Brazil (Fontana et al. 2008; Bencke et al. 2010), and to neighboring countries – Uruguay (Aldabe et al. 2009) and Argentina (Chebez et al 2011). Globally, this species is included within the “near-threatened” category and its populations have been decreasing across its distributional range (BirdLife International, 2016). At national scales, the scarce knowledge about its population size and life history, along with its reduced distribution and high habitat specificity has led to its inclusion in threatened species lists in Uruguay and Argentina, under “near-threatened” (Aldabe et al. 2009) and “threatened” (Chebez et al. 2011) categories, respectively. In Brazil, the Straight-billed Reedhaunter has been recently excluded from the list of threatened species (MMA 2014)

*Gravatazais* present high biological, taxonomic and functional diversity, including the presence of many other threatened species, such as annual fishes, amphibians, birds and mammals (Fontana et al. 2003, Teixeira de Mello et al. 2011; Lanés et al. 2014). However, in southern Brazil, regions with high concentration of this type of wetland are also characterized by low annual precipitation, and the constructions of dams have been very important for the development of cities and local communities (Boschi et al. 2011). This is of particular concern as dams significantly affect the population dynamic of terrestrial species (Kingsford 2000).

In this work, we aimed to identify the effects of habitat fragmentation on the population dynamics of the Straight-billed Reedhaunter. Specifically, we determined the probabilities of occupation, colonization, and extinction associated with the habitat changes promoted by the construction of a water reservoir. For this purpose, *gravatazais* with and without Straight-billed Reedhaunter subpopulations were monitored for their presence and absence during the reservoir construction activities. Given the close relationship of Straight-billed Reedhaunter to wetlands dominated by Eryngo, we expected to find significant effects of habitat fragmentation on the persistence of their subpopulations, which would reflect on the extinction and colonization probabilities.

## Methods

### Study area

The study was conducted in the municipality of Bagé in southernmost Brazil (S 31° 17’ 26,4’ and W 54° 09’ 32,7’). This region greatly represents the landscape of the Pampa Biome (Rambo 1959; Overbeck et al. 2007). The study area is characterized by large extensions of natural grassland, marked by the constant presence of livestock (main economic activity in the region) and agricultural crops, such as corn, soy, sorghum and exotic forest (IBGE 2013). Specifically, the place designated for the reservoir construction covers an area of approximately 340 hectares (Figure 1). Inside the construction area, an extensive riparian forest surrounds the most important watercourse of the locality, called “Arvorezinha”. Grasslands and drainage lines with grass and low shrubs are the dominant cover inside the area, as well as patches of wetlands dominated by sedge (*Eryngium pandanifolium*), reed (*Cyperus californicus*), cattail (*Typha dominguensis*) and panic grass (*Panicum prionitis*).

**Fig. 1.**
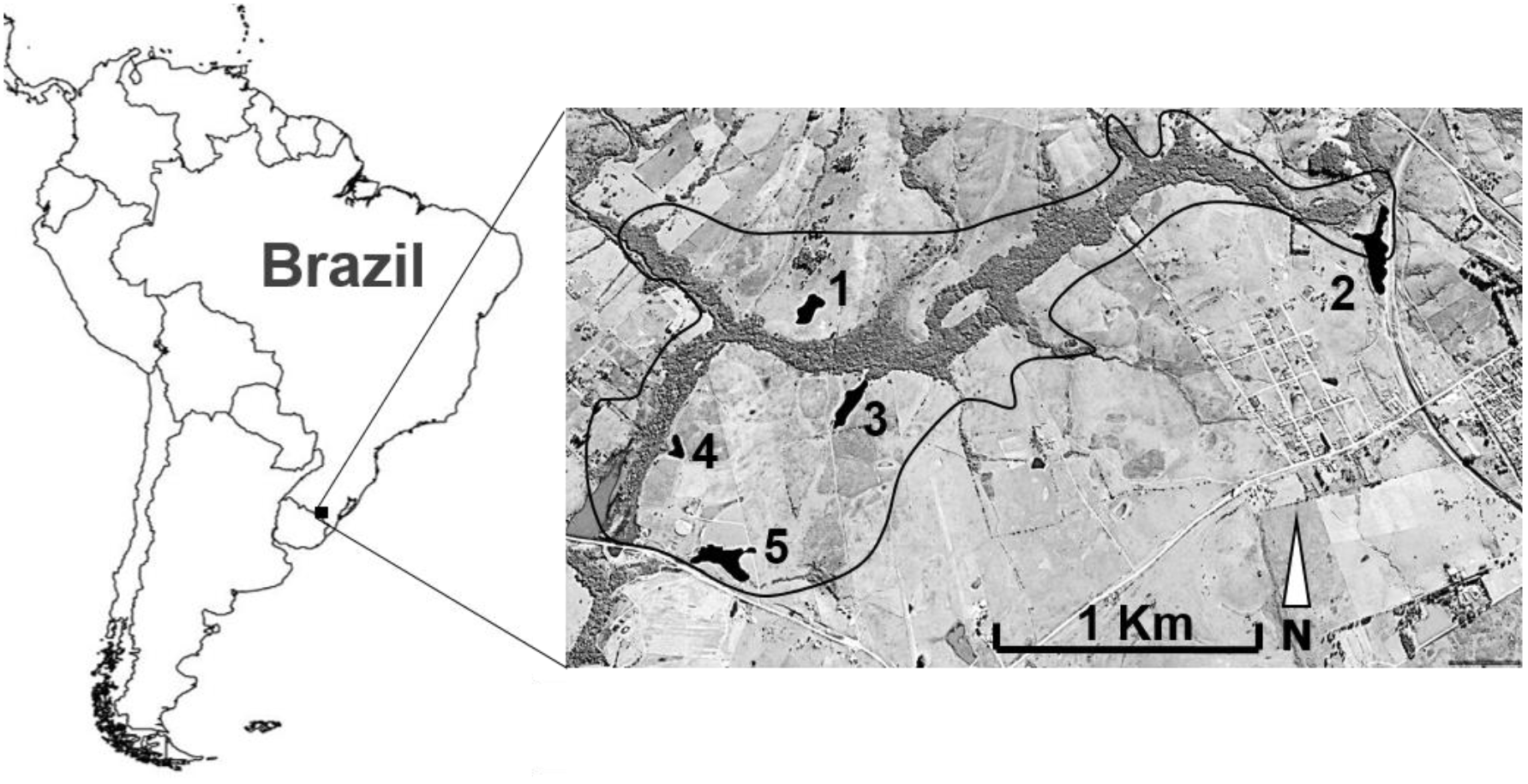
Study area located in southernmost Brazil. The continuous line demarcates the approximated flooding area of the reservoir. Selected patches are numbered and marked. Image dates to April 2007 taken from Google Earth Pro (accessed on 12 June 2016).

### Patch selection and population monitoring

Firstly, from satellite imagery, we identified all potential wetlands within the study area. During a pilot inspection in December 2011, we excluded all sites without Eryngo habitat. Five Eryngo wetlands patches were selected for research (Table 1). The size of wetlands varied between 5444 m^2^ and 34150 m^2^. The presence of two individuals of Straight-billed Reedhaunter was confirmed in three of these patches, during the pilot study. Posteriorly, between September 2012 and April 2013 (except for November), we conducted seven monthly visits to the five selected wetlands. In this sense, including the first observation performed during the pilot inspection, eight sampling visits were performed for each wetland. Nests were recorded in three patches initially occupied by the Straight-billed Reedhaunter – see Gonçalves et al. (2017) for more information about breeding biology.

**Table 1.**
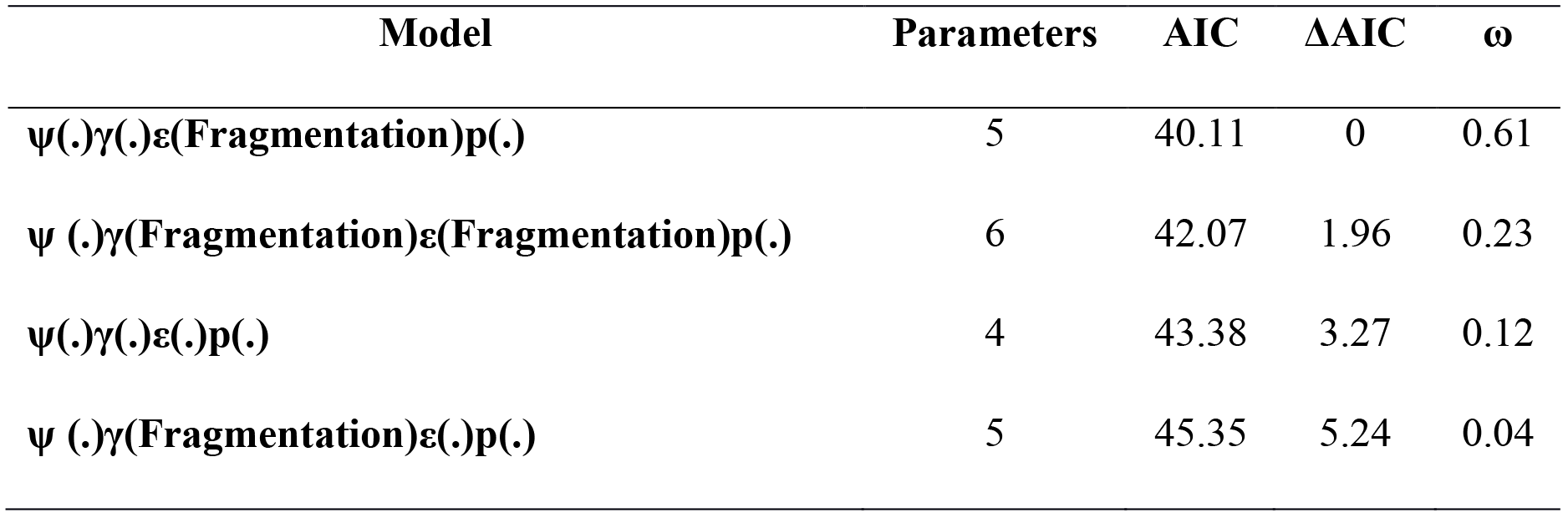
Model-selection table with candidate models ranked according to their AIC weights(ω), from highest values to lowest values.

During each visit, we aimed to detect the presence/absence of Straight-billed Reedhaunter and whether wetlands were fragmented. We considered fragmentation to be any change in the natural structure of a wetland that implied a new split of the continuous patch. All samplings were made when weather conditions were favorable (without rain and little wind). To facilitate the detection of Straight-billed Reedhaunte, observations were always made by two researchers. “Playback” techniques were also run on the edge of patches and the search time in each site varied between 1 and 2 hours.

### Occupancy models

The presence and absence of Straight-billed Reedhaunter was modeled as a function of habitat integrity (occurrence or not of one event of fragmentation or destruction of the wetlands). Four parameters were evaluated: probability of occupation (ψ); probability of colonization (*γ*); probability of extinction (*ε*) and probability of detection (p) (this last was kept constant in all models as the species can be easily detected). We tested the fit of four models: 1) ψ(.)*γ*(.)ε(.)p(.) – model in which the probability of occupation, colonization, extinction and detection are constants (.); 2) ψ(.)γ(fragmentation)ε(fragmentation)p(.) – model in which fragmentation affects colonization and extinction probabilities; 3) ψ(.)γ(fragmentation)ε(.)p(.) – model in which fragmentation affects only the probability of colonization; and 4) ψ(.)γ(.)ε(fragmentation)p(.) – model in which fragmentation affects only the extinction probability. We evaluated model fit using a multimodel inference approach within an information-theoretic framework (Burnham & Anderson 2003, Anderson 2008). To estimate model plausibility we used Akaike Information Criterion (AIC) and AIC weight (*w*), which measures the relative likelihood of the model given the data, normalized across the set of candidate models (Burnham & Anderson 2003, Anderson 2008). Occupancy models were fitted using the Unmarked Package (Fisk and Chandler 2015), in the R v3.1.3 environment (R Development Core Team 2014).

## Results

The spatial-temporal dynamic of the Straight-billed Reedhaunter is presented in Figure 2. Of these three wetlands, two were not altered by the reservoir construction and the presence of individuals was constant throughout the study period. One of the occupied patches (number 3, Figure 2) was partially destroyed during the breeding period (Figure 3). During the next sampling periods, the species was absent in this patch, and individuals were concomitantly observed in two other initially unoccupied wetlands, including one already fragmented site (Fig. 2a-2d).

**Fig. 2.**
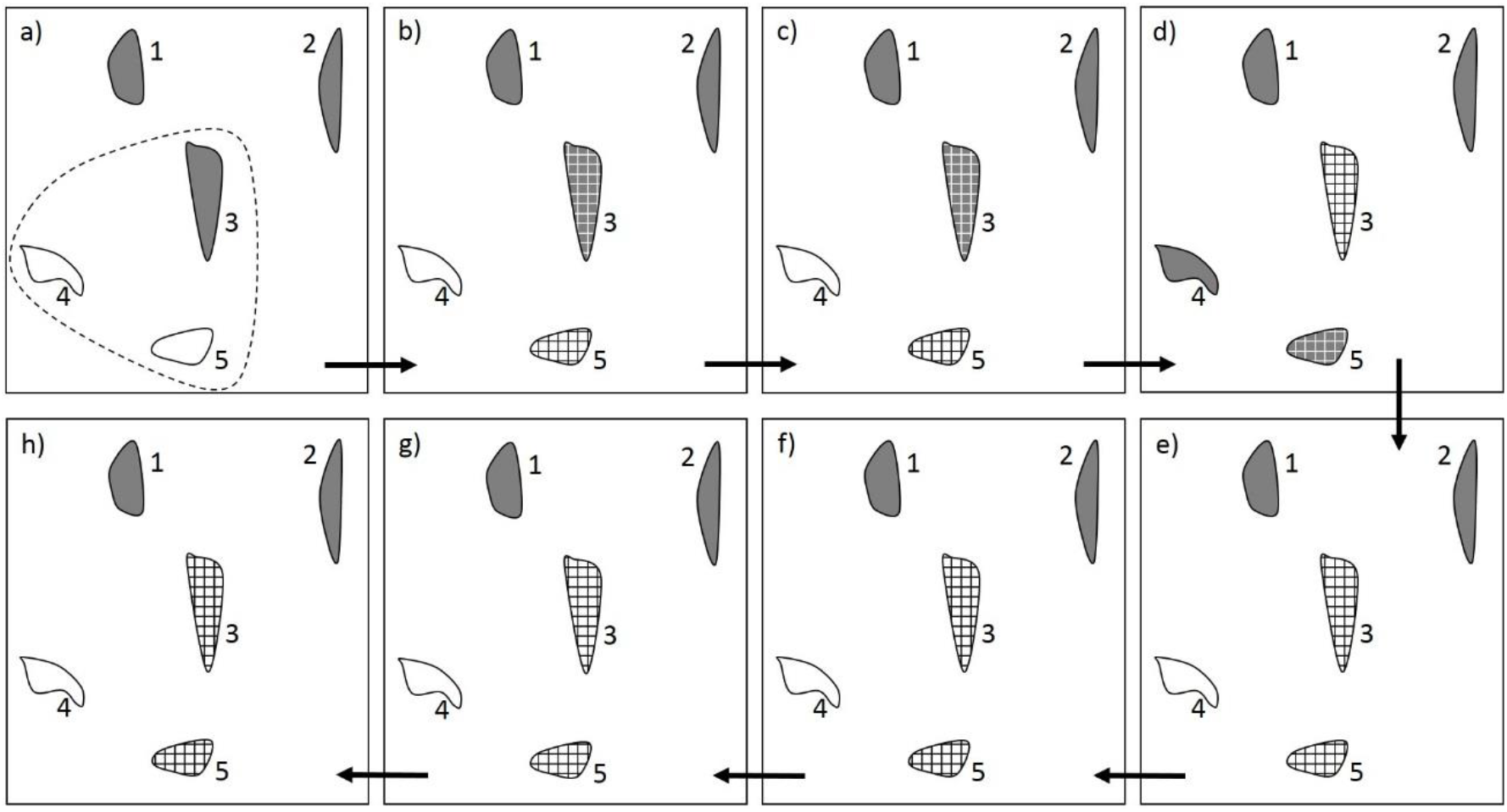
Schematic representation of the Straight-billed Reedhaunter’s occupation dynamics. A dotted line in Figure 2a marks the area affected by the construction activities. The areas with the presence and absence of species are filled in gray and white, respectively. The patches with gridlines represent the occurrence of a fragmentation event. The dates of the samples were: 2a) December 2011; 2b) September 2012; 2c) October 2012; 2d) December 2012; 2e) January 2013; 2f) February 2013; 2g) March 2013; and 2h) April 2013.

**Fig. 3a-3c.**
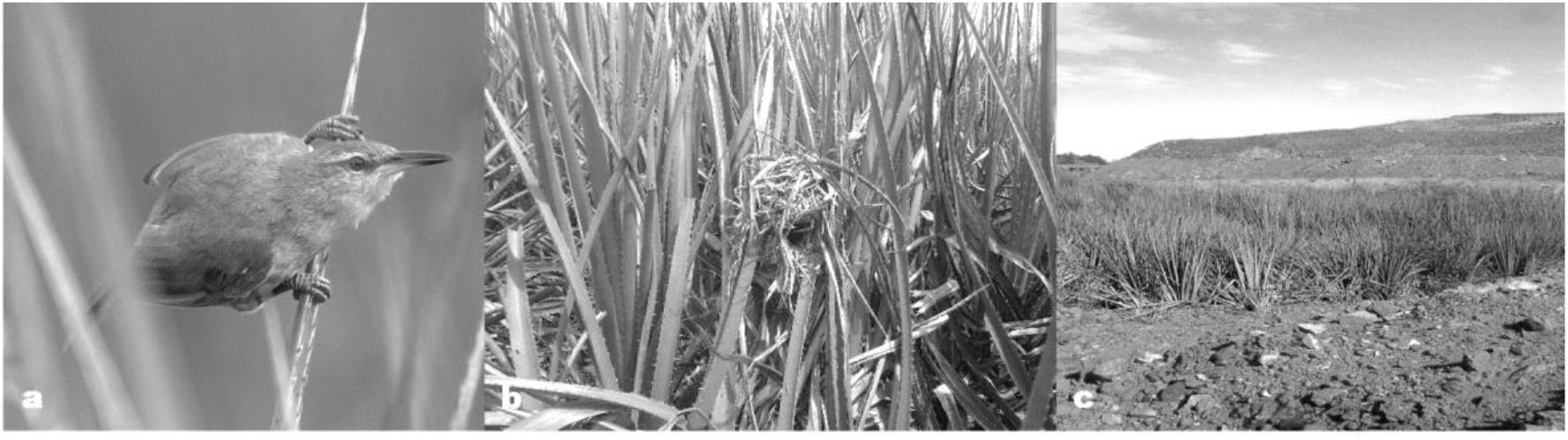
3a,Straight-billed Reedhaunter perched on *Eryngium pandanifolium;* 3b, Nest built on the stems of “Eryngo”; 3c, wetland partially destroyed by the reservoir construction activities. Photos: 3a, Christian Andretti; 3b and 3c, Priscila Pons.

The model in which fragmentation affected only the probability extinction was the most plausible (Tab. 1). Model in which fragmentation affects both the colonization and extinction was also highly plausible (ΔAIC < 2), although its probability was much lower than the model in which the probability of extinction was a function of habitat alteration (Tab. 1). Extinction probability of Straight-billed Reedhaunter tended to increase drastically in fragmented wetlands (Tab. 2; Fig. 4).

**Fig. 4.**
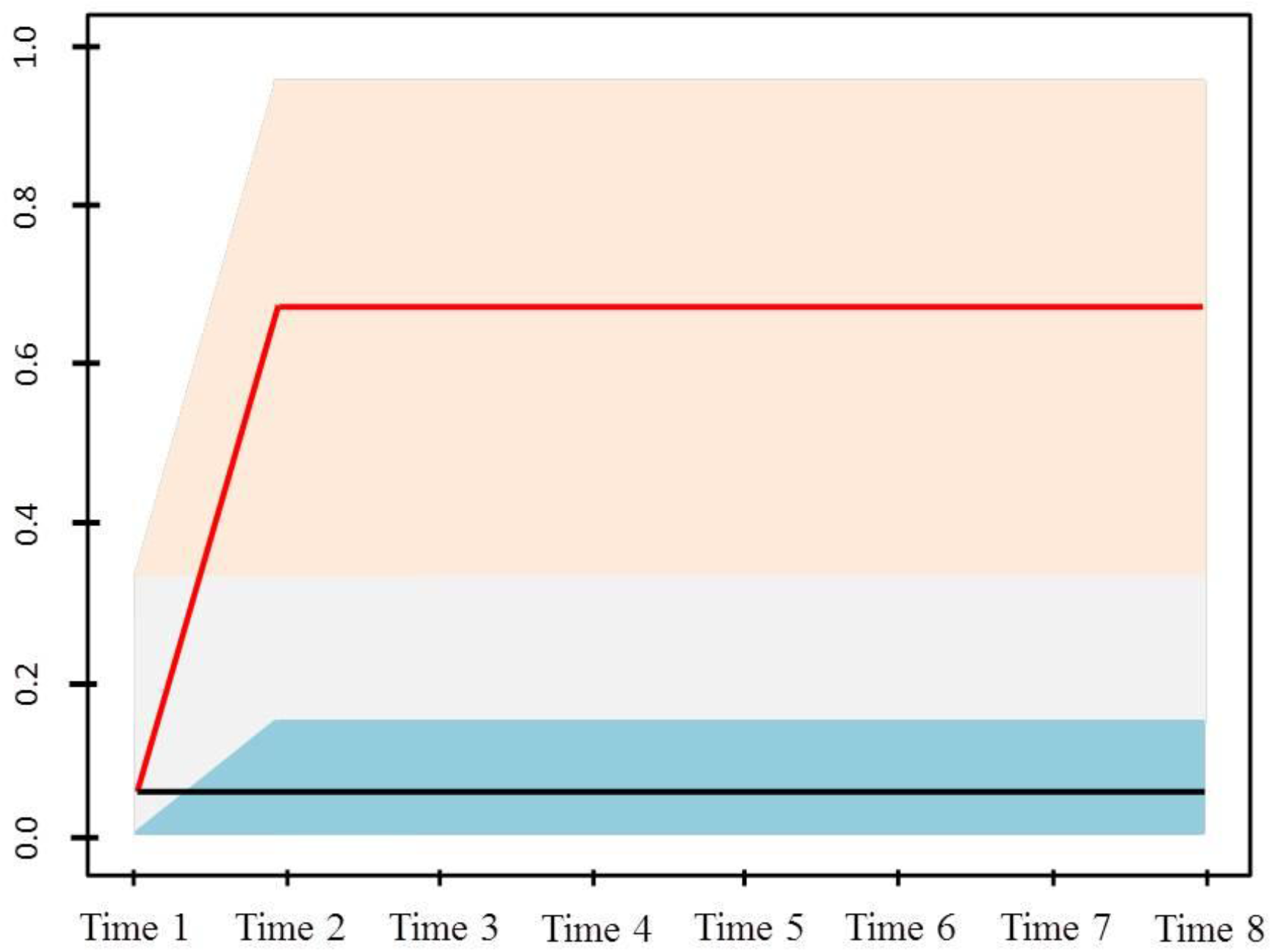
Extinction probability (95% Confidence Interval) of Straight-billed Reedhaunter estimated foreach sampling month in areas with and without fragmentation. Orange: fragmented patches; Grey: non-fragmented patches.

**Table 2.**
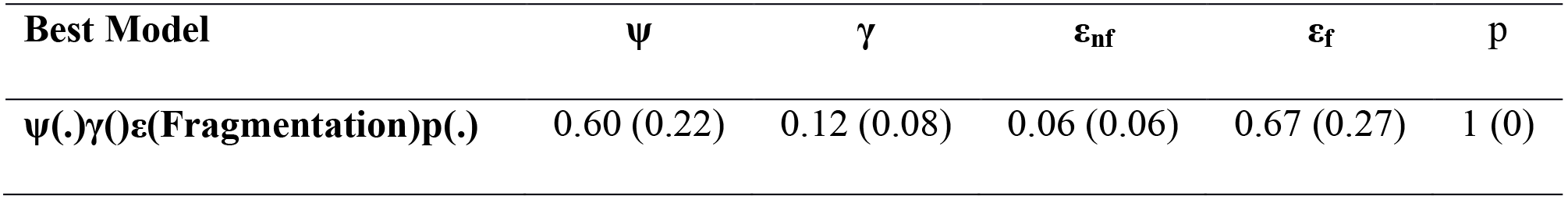
Probabilities estimates (±se) for the Straight-billed Reedhaunter for the best model after the fragmentation event. Subscript for extinction probabilities denote non-fragmented (ε_nf_) and fragmented patches (ε_f_).

## Discussion

Our results highlight that the effect of habitat fragmentation on Straight-billed Reedhaunter populations is significant. Fragmentation increases the extinction probability of the Straight-billed Reedhaunter.

Habitat fragmentation results primarily in local extinction of populations with consequences to their patterns of regional and global distribution (Henle et al. 2004). Although species are able to occupy fragmented landscapes when their life cycles include multiple fragments (Redpath 1995), the effects of fragmentation is stronger in those species with specific ecological requirements (Swihart et al. 2003; DeVictor et al. 2008). This seems to be the case of the Straight-billed Reedhaunter. After local extinction in one of the wetlands, individuals were sporadically observed in two other initially unoccupied patches, including a fragmented site. Although we cannot confirm that these individuals are the same as those in the previously destroyed wetlands, the temporary occurrence of the Straight-billed Reedhaunter in these sites could indicate an immediate attempt to extend its territory, as a result of the habitat loss and displacement of competitive individuals.

The size and stability of the populations in natural wetlands depend strongly on a range of habitat and landscape factors, as well as on body size, morphology, behavior, effects of niche breadth and the effect of geographic range boundaries (Redpath, 1995; Marsh and Trenham 2000; Swihart et al. 2003). Stable subpopulations were observed only in unaltered wetlands, which indicate that the species’ ecological plasticity is apparently low. Paradoxically, the species has been known to occupy wetlands in widely altered landscapes, e.g., on the edge of roads and dams (Ricci and Ricci 1984; Barbarskas and Fraga 1998). We recommend taking these observations cautiously as the populations may have colonized these wetlands after the alteration of the landscape, which favors the colonization of Eryngo due to artificial water concentrations by construction of roads and dams. Hence, further research is needed to elucidate the effects of the area and habitat structure on the size of the subpopulations located in unchanged natural patches.

Our results show the Straight-billed Reedhaunter populations have greater stability in unaltered patches, which reduces the chances of permanence and colonization in those patches subjected to fragmentation events. These results demonstrate a more accurate view of this species’ ecological plasticity and their tolerance to habitat fragmentation, and contribute to lower the uncertainties of its degree of threat at different scales. The Straight-billed Reedhaunter is globally near-threatened (BirdLife International, 2016), but has been recently excluded from the list of endangered species in Brazil (MMA, 2014) and Rio Grande do Sul State (Rio Grande do Sul, 2014). Indeed, the number of records of this species has significantly increased in recent years, especially in southernmost Brazil (Develey et al. 2008; ICMBIO, 2014; Wikiaves, 2016). On the other hand, it is important to note that the knowledge about many aspects of its biology and ecology is completely lacking. In this work, we discuss the effect of fragmentation promoted by an activity that tends to drastically alter landscapes. Additionally, these wetlands are commonly channeled for water subtraction – a practice that also fragments wetlands and may have similar consequences to the disturbance explored herein. Therefore, we strongly recommend understanding the relationship of both the habitat structure and degree of connectivity of patches with the Straight-billed Reedhaunter’s spatial distribution by identifying how environmental and spatial stochastic processes on different landscape scales influence the species demographic dynamics. Finally, the natural distribution of the *gravatazais* may imply that the Straight-billed Reedhaunter may be distributed as a metapopulation, and we end this article by encouraging further research about not only the Straight-billed Reedhaunter, but also about the vast biodiversity associated with wetlands dominated by Eryngo.

## Acknowledgements

We are grateful to the Ecossis Soluções Ambientais for support while undertaking fieldwork.

